# Impact of microchannel width on axons for brain-on-chip applications

**DOI:** 10.1101/2024.05.16.594497

**Authors:** Katarina Vulić, Giulia Amos, Tobias Ruff, Revan Kasm, Stephan J. Ihle, Jöel Küchler, János Vörös, Sean Weaver

**Affiliations:** Laboratory of Biosensors and Bioelectronics (LBB), Gloriastrasse 37/39, Zürich, Switzerland

## Abstract

Technologies for axon guidance for *in vitro* disease models and bottom up investigations are increasingly being used in neuroscience research. One of the most prevalent patterning methods is using polydimethylsiloxane (PDMS) microstructures due to compatibility with microscopy and electrophysiology which enables systematic tracking of axon development with precision and efficiency. Previous investigations of these guidance platforms have noted axons tend to follow edges and avoid sharp turns; however, the specific impact of spatial constraints remains only partially explored. We investigated the influence of microchannel width beyond a constriction point, as well as the number of available microchannels, on axon growth dynamics. Further, by manipulating the size of micron/submicron-sized PDMS tunnels we investigated the space restriction that prevents growth cone penetration showing that restrictions smaller than 350nm were sufficient to exclude axons. This research offers insights into the interplay of spatial constraints, axon development, and neural behavior. The findings are important for designing *in vitro* platforms and *in vivo* neural interfaces for both fundamental neuroscience and translational applications in rapidly evolving neural implant technologies.

## 1 Introduction

Neurons are the building blocks of the nervous system having a unique structure and polarity [1, 2, 3] essential for efficient information transmission. The polarity of neurons is expressed by two distinct subcellular components: dendrites and axons. Dendrites are projections emerging from one side of the cell soma, acting as the main computational units of the neuron [4]. They receive information from multiple presynaptic partners through synapses which can trigger action potentials (AP) to propagate through axons. AP elicitation and subsequent propagation are the primary form of communication between neurons [5, 6]. Axon growth is vital for functional development and potential repair of the nervous system. It relies on an interplay between extracellular and intracellular cues [7], whose dynamics are still not fully understood. Given the challenges studying the complex central nervous system *in vivo* [8, 9, 10, 11], isolated mechanisms of axon growth have been extensively studied *in vitro*[1, 12, 13, 14, 15]. A substantial amount of research has been done with rat primary cortical [16, 11], thalamic [17, 11], and hippocampal [18] neurons. For these cells, several stages of growth and development have been studied and defined. Within hours of seeding, neurons undergo morphological changes from rounded symmetrical cells to polar cells with distinct dendrites and axons. Over the next days *in vitro*, the axon growth cone, a mobile and flexible sensory structure at the axon tip, explores its environment while receiving guidance cues from the extracellular matrix and neighboring axons to enforce the direction and extent of axon growth [19]. The tension forces between an axon and a substrate [20], and forces between axons (axon fasciculation [21]) play an important role in determining their growth extent and trajectory. Understanding and leveraging these mechanisms is crucial not only for understanding nervous system development but also for exploring the potential of axon regeneration for advancing therapeutic interventions in cases of nerve damage.

Studying neurite outgrowth in random cultures hinders our understanding of the development of individual axons and the factors governing axon length. For instance, distinguishing axons from axon bundles is challenging due to inherent axon size variability [17, 22] hampering the tracking of unique developmental trajectories. Differences in development between isolated axons and axon bundles further complicate the analysis. Additionally, assessing the impact of extracellular cues on growth cone exploration or accurately measuring axon length face complications in the absence of a controlled environment. To establish more controllable and reproducible experimental systems, researchers have increasingly turned to controlling neural network topology through patterning. Some methods applied to control the network topology *in vitro* have been inspired by *in vivo* studies that demonstrate directional guidance through adhesion forces [23]. The aforementioned interactions of axons with a substrate and axon fasciculation are both critical to consider in the design of culture tools for axon guidance. By providing controlled surfaces with cell-adhesive coatings or adjusting the mechanical properties of substrates, researchers can guide axon growth and explore the interplay between neurons and their environment [24]. The substrate can be modified by techniques such as microcontact printing,[16, 20] photolithography [25, 26], electrochemical surface modification [27] and light-induced nanotopography [28] in order to manipulate network topology. Notably, patterning methodologies can be combined with microelectrode arrays (MEAs)[29, 30, 27], allowing for the recording and modulation of extracellular neural activity, thus providing a functional aspect to create engineered neural systems. This way neural development can also be monitored in terms of functionality in parallel with its morphology.

Despite the benefits from the described patterning methods, there remain challenges to be addressed. Coatings may degrade over time or be influenced by axon forces, limiting their suitability for extended culturing [24, 31, 32]. Achieving precise neurite guidance [33] and controlling neural network size [24, 34] remains challenging. To address these concerns, an alternate patterning approach involves physical confinement through polydimethylsiloxane (PDMS) microstructures. PDMS microstructures are biocompatible and gas-permeable, hence they support long-term cell culturing [34]. Their transparency facilitates high-resolution imaging, and compatibility with MEAs allows for integrated functional analysis [35]. The diverse design possibilities of microfabricating PDMS enable subcellular compartmentalization and precise control over directionality [36, 35, 37, 38]. Researchers can tailor the network size and topology to address specific phenomena, providing a versatile platform for exploring neural dynamics.

In the context of axon development, PDMS microstructures offer an opportunity to investigate spatial limitations of axon growth. It has been observed axons tend to follow edges and avoid turning in more than 90 degree angle [39, 40, 41]. The microchannel height influence on electrophysiology recording fidelity has been systematically addressed using two-compartment microfluidic systems.[42] However, the effects of spatial constraints in terms of width of the microchannels on axon growth, specifically in the lower, submicron range has not yet been studied in detail. For example, it is still unknown how constraints affect the extent of axon bundle growth, how sensitive conduction speed of axon bundles is to axon bundle size and if spatial constraint modifies the speed. Furthermore, growth cone adaptability to spatial constraints is not fully explored. How they regulate their length accordingly to their width, for instance on a surface considered as effectively infinite along one direction, has not been quantitatively addressed in detail [43]. The morphological adaptability of growth cones, spanning dimensions from a few microns to several tens of microns,[1, 44, 45, 46] empowers axons to dynamically adjust to the diverse extracellular spaces they encounter. This adaptability is particularly important in navigating divergent extracellular environments found *in vivo*.

In this work we investigate how variations in the number of efferent channels and channel size of PDMS microstructures impacts axon growth rate, bundle formation, and activity propagation. By reducing the size of the topological restriction to the same order or smaller than the growth cone we study the narrowest space restriction axons can penetrate.

## 2 Materials and Methods

### 2.1 PDMS microstructures

PDMS microstructures for cell and axon guidance were designed in a CAD software (AutoCAD 2021) and fabricated by Wunderlichips (Zurich, Switzerland). The fabrication process is described in former publications [41, 36]. All microstructures used in this study have wells where the cells are seeded and microchannels which are impermeable for soma but accessible for neurites. Schematics of the microstructures are shown in Fig.1. The PDMS thickness and microchannel height vary between structure types. 200 *µ*m thick microstructures will be referred to as spheroid-seeding PDMS microstructures and 75 *µ*m thick microstructures as cell-suspension-seeding PDMS microstructures.

**Fig. 1:**
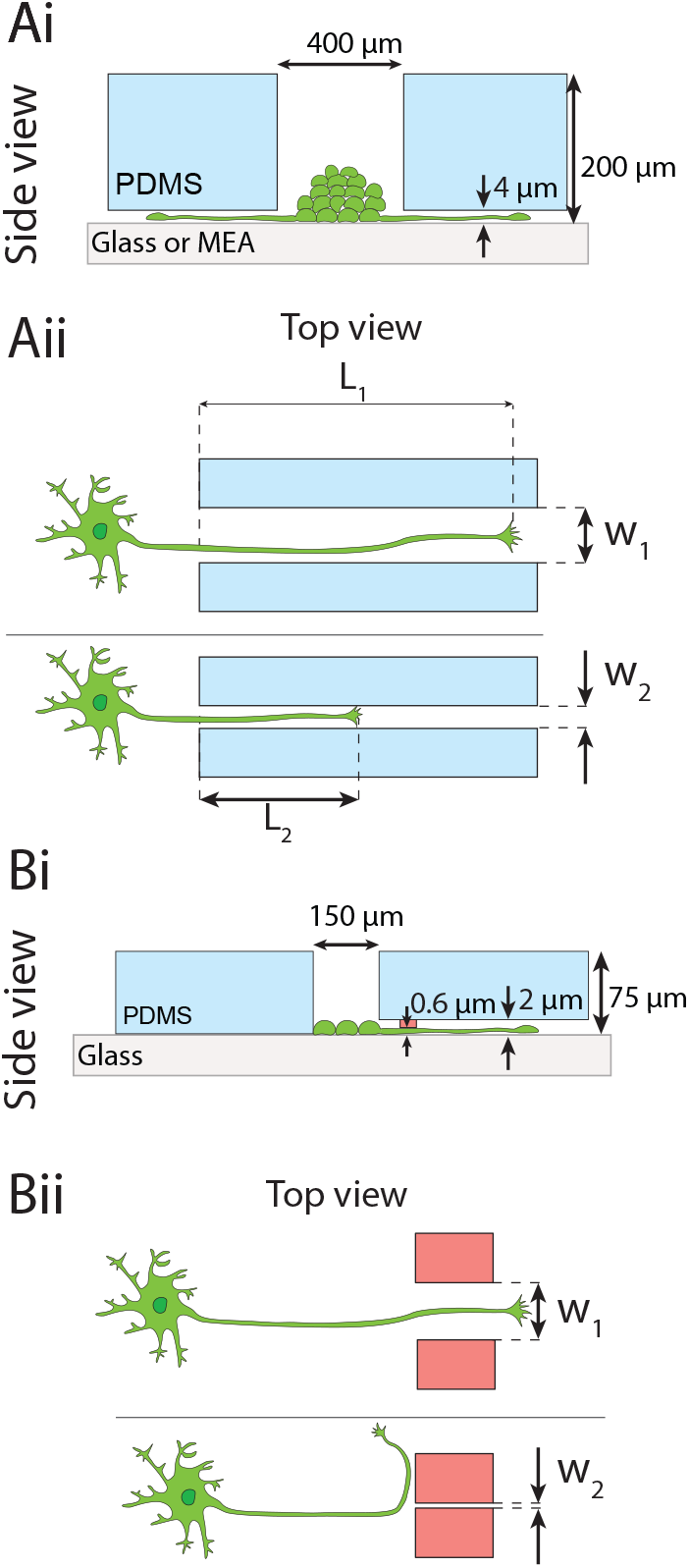
Overview of the PDMS microstructures used in this study for spatially confining axon growth. Ai) Side view of the spheroid-seeding microstructure. Spheroids are seeded into the wells. Microchannels that emerge from the wells are too narrow for the cell soma to grow through but allow for axon growth. Aii) The length of axons, starting from the beginning of the microchannel, was investigated for different microchannel widths. Bi) Side view of the thin microstructure. Low-density cell suspension is seeded into the wells. Microchannels contain additional submicron constraint (depicted in red). Bii) The spatial limitation of axon growth is assessed by noting what is the smallest constraint a growth cone can penetrate.

#### 2.1.1 Spheroid-seeding PDMS microstructures

The PDMS microstructures used in the experiments investigating the effect of channel width in Fig.2 and Fig.3 are 200 *µ*m thick and consist of a 400 *µ*m diameter well to accommodate neural spheroids. The well then branches into microchannels. Each microchannel is 4 *µ*m high along the whole length. The 4 *µ*m height of the microchannels makes it impermeable for cell soma but accessible for axons (and dendrites). Microchannels are 8 mm long. The PDMS microstructure shown in Fig.2 have 44 concentric microchannels emerging from the central seeding well that narrow down to 1.5 *µ*m in width to provide similar starting conditions. After the 1.5 *µ*m wide constriction, these channels branch into microchannels with widths varying from 1.5 to 75 *µ*m. The design has a point symmetry so each half is considered as a separate replicate. The PDMS microstructure shown in Fig.3 has microchannels with the identical width of 50 *µ*m.

**Fig. 2:**
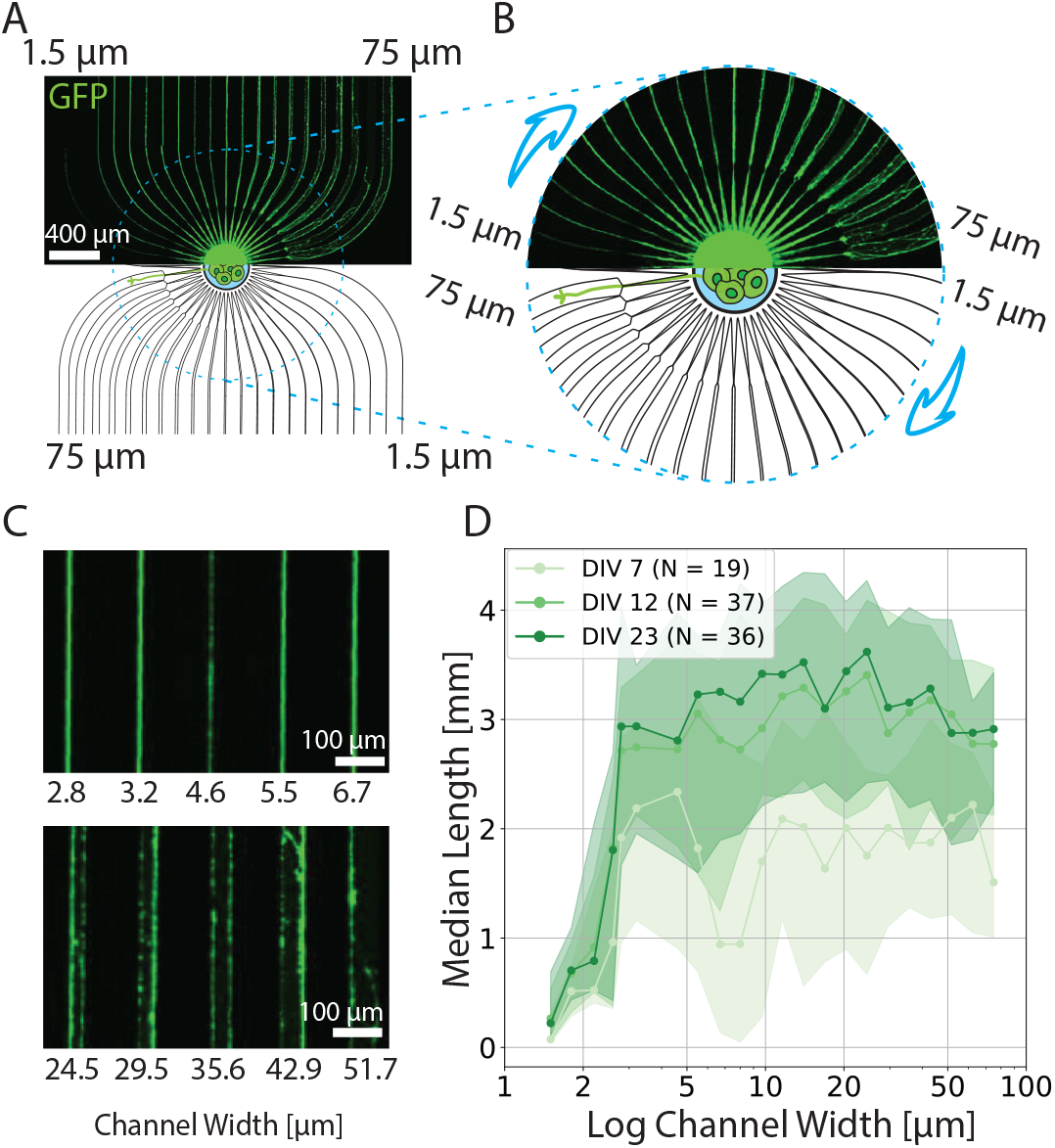
Spheroid-seeding PDMS microstructure for studying the length of axons for different microchannel widths. A) Overview of the microstructure. The bottom half shows the schematic of the microstructure, while the upper half shows neurons expressing GFP at DIV 12. B) Close up image of the seeding well and microchannels emerging from it. All microchannels initially narrow down to 1.5 *µ*m in width and then extend to the final microchannel size that increases from 1.5 to 75 *µ*m as indicated by an arrow. C) Example of axon growth through various microchannel widths. D) Median axon length with the corresponding quartile range indicated as shading as a function of channel width for different DIV. Note that x axis is represented by a logarithmic scale. Number of biological replicates is 6.

**Fig. 3:**
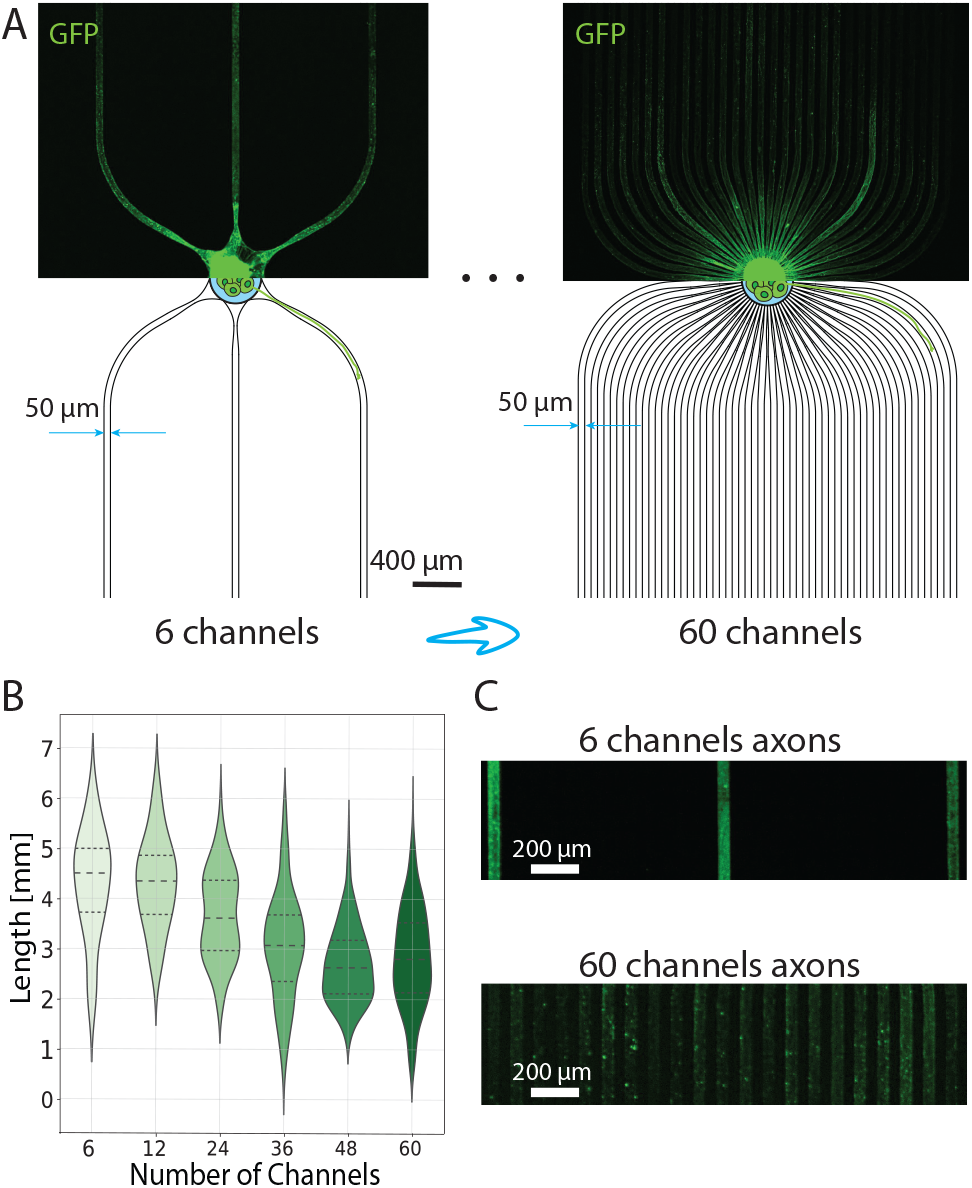
Controlling the number of axons per microchannels with spheroid-seeding PDMS microstructures with different number of microchannels emerging from the central well. A) The two extreme examples of the microstructures. The bottom half shows the schematics and the dimensions of the microstructures, while the upper half shows GFP-expressing neurons at DIV 23. The number of microchannels was varied from 6 to 60. B) Example of axons growing in the 6-channel (top) and 60-channel(bottom) microstructure. C) Violin plot with a corresponding quartile range for the axon length dependency on the number of microchannels emerging from the central well. The number of technical replicates is 7, 8, 10, 8, 8, 7 in the ascending order of the microchannel number.

#### 2.1.2 Cell-suspension-seeding microstructures with variable submicron tunnel width

The thickness of the PDMS microstructures described in Fig5,6 is 75 *µ*m and it consists of eight wells of area 120 x 260 *µm*^2^ that are suitable for seeding cells in suspension. They feature 2 *µ*m high microchannels with submicron size, 600 nm high tunnels. In the case of microstructures shown in Fig.5, seven seeding wells narrow down to nano/submicrochannel tunnels with a height of 600 nm and width varying from 0.6 to 1.8 *µ*m which then extend to a width of 30 *µ*m. The eighth channel has a fixed width of 30*µ*m and serves as a control. The PDMS microstructure described in Fig.6 consists of a hexagonal seeding well with a 145 *µ*m long side that narrows down to thirteen tunnels with widths varying from 150 to 1000 nm. The microchannels that follow are 0.65 mm long.

### 2.2 Substrate preparation

#### 2.2.1 Glass bottom dish

Culture dishes with microscopy glass bottom were used for imaging experiments. 30 mm diameter glass coverslips (Menzel glass, selected no. 1.5, ThermoFisher) were mounted to plastic rings (WillCo Wells) according to the manufacturer’s instructions. Mounted dishes were filled with isopropanol, ultrasonicated for 10 mins and afterwards further rinsed with isopropanol, and ultrapure water (Milli-Q, Merck-MilliPore) before being blow-dried with nitrogen.

Next, the dishes were treated with air plasma for 2 min (18 W PDC-32G, Harrick Plasma) and coated with 500 *µ*L per dish of 0.1 mg mL^−1^ Poly-D-lysine (PDL) (P6407, Sigma Aldrich) in Phosphate buffered saline (PBS) (10010-023, ThermoFisher). After PDL coating at room temperature for 45 min, dishes were rinsed three times with ultrapure water. Ultrapure water was then aspirated and the dish was carefully blow-dried with nitrogen.

#### 2.2.2 Microelectrode array

Glass MEAs (60MEA500/30iR-Ti-gr, Multichannel Systems) with a 6×10 electrode grid, electrode spacing of 500 *µ*m, and an electrode diameter of 30 *µ*m were used for experiments requiring extracellular activity recordings. MEAs were reused across several experiments. When reused, MEAs were rinsed with ultrapure water and immersed in 4 % Tergazyme (1304-1, Alconox) overnight to remove organic debris. MEAs were immersed in ultrapure water and placed in the fridge for long-term storage. Prior to cell seeding, MEAs were rinsed with isopropanol and ultrapure water and dried with nitrogen. After thorough drying, MEAs were treated with air plasma for 2 min and coated with 500 *µ*L per MEA of 0.1 mg mL^−1^ PDL for 45 min at room temperature. MEAs were afterwards rinsed with ultrapure water and carefully blow-dried with nitrogen.

#### 2.2.3 Microstructure Attachment

The PDMS membrane was cut with a scalpel and an individual microstructure was placed on the substrate with tweezers. For MEAs, the microchannels were aligned along the electrodes under a light microscope (Leica Microsystems, Germany) using a drop of ultrapure water and tweezers. Upon placing the microstructure on the substrate and subsequent visual inspection the dishes were left in the oven at 37 ^*°*^C for 45 min. Afterwards, PBS was added in the dishes and they were placed in the desiccator for 10 min or until there was no air bubbles coming out of the microstructures. After desiccation, PBS was exchanged with NeuroBasal (NB) medium and the dishes were ready for cell seeding.

### 2.3 Cell culture

The medium used for culturing the cells is NeuroBasal medium (NB) (21203-049, ThermoFisher) [47]. NB complete medium was prepared freshly. NB complete medium is 2 % solution of B-27 supplement (17504-044), 1 % solution of Penicillin-Streptomycin (P-S) (15070-063) and 1 % solution of GlutaMAX (35050-061, all from ThermoFisher).

#### 2.3.1 Cell dissociation

Primary thalamic or cortical neurons from E18 embryos of pregnant Sprague-Dawley rats (EPIC, ETH Phenomics center) were used in the experiments. Animal experiments were approved by the Cantonal Veterinary Office Zurich. Embryonic neuronal tissue was dissected and stored in hibernate E medium (ThermoFisher A1247601) on ice. Cell dissociation began by digesting the tissue in a solution consisting of 50 mg mL^−1^ Bovine serum albumin (BSA) (A7906, Sigma-Aldrich), 1.8 mg mL^−1^ D-glucose (Y0001745, Sigma-Aldrich), and 0.5 mg mL^−1^ papain (P5306, Sigma-Aldrich) dissolved in sterile PBS. Directly prior to dissociation, the solution was warmed to 37 ^*°*^C, filtered (0.2 *µ*m) and 1 mg mL^−1^ DNAse (D5025, Sigma-Aldrich) was added. Tissue was left in the papain solution for 15 minutes at 37 ^*°*^C, after which the solution was replaced by NB medium with 10% fetal bovine serum (10500056, ThermoFisher) to stop the digestion. Two subsequent washes with NB were done, waiting 5 minutes between each wash. This was followed by trituration and cell counting (Cell Countess, Invitrogen). Cells dissociated from a single pregnant rat were considered one biological replicate.

#### 2.3.2 Cell seeding

Cells were seeded either as spheroids or in suspension, depending on the experiment.

#### 2.3.3 Spheroid preparation and seeding

In the case of microstructures described in section 2.1.1, cells were seeded as spheroids. To prepare the spheroids, after counting the cells, the volume of cell suspension needed to create 500-cell spheroids was added to an AggreWell microwell plate (Stemcell Technologies Inc., Canada). The wells were then filled with NB to a volume of ∼2 mL. The cells were labeled with enhanced green fluorescent protein (eGFP) in the AggreWell plate by adding ∼10k particles/cell of the adeno-associated viral (AAV) vector (V-DJ/2-hSyn1-chl-EGFP-SV40p(A) (University of Zurich Viral Vector Facility). After transduction, the AggreWell containing the cells was centrifuged for 3 min to ensure homogeneous cell distribution and formation of spheroids. The spheroids in the AggreWell were kept in the incubator until seeding.

The day after dissociation, spheroids formed in the AggreWells were ready for seeding in the PDMS microstructures. For seeding, 0.5 mL of spheroid solution was transferred to a small Petri dish to avoid leaving the whole AggreWell at room temperature and without CO_2_. 5-10 spheroids were aspirated from a Petri dish using a 10 *µ*L pipette and a spheroid was carefully pipetted into the center wells of the microstructures one-by-one. Once all seeding spots were filled but no longer than 10 min after the spheroids were removed from the incubator, the culture (a glass dish or a MEA) was placed in the incubator at 37 ^*°*^C, 90% humidity and 5% CO_2_. Every three to four days, ∼0.5 mL of the old medium was exchanged with ∼0.6 mL of the fresh medium. In the experiments with spheroids, the day when spheroids were seeded in the PDMS microstructures was considered day in vitro (DIV) zero.

#### 2.3.4 Cell suspension seeding

In the case of 75 *µ*m-high microstructures, larger spheroids were too large for seeding into the wells without falling out due to the smaller PDMS thickness and smaller spheroids were too small for manual seeding. Hence, neurons were seeded in suspension. After counting the cells in the solution upon dissociation, the exact volume to obtain roughly 21000 cells/mm^2^ was calculated. Cells were transduced with eGFP AAV and mRuby3 ((scAAV-DJ/2-hSyn1-chl-mRuby3-SV40p(A)) AAV. The cells were seeded with a pipette centered at the top of the PDMS microstructures to increase the probability of cells falling inside the wells. 20 min upon seeding, dishes were inspected under the microscope to assess if enough cells have fallen inside the seeding wells. In case there was not enough neurons in the wells, they were re-suspended by pipetting the medium up and down above the PDMS microstructure. In the experiments with such cell suspensions, day of cell dissociation and seeding was considered DIV zero.

### 2.4 Image acquisition and analysis

Transduced cultures were imaged using a confocal laser scanning microscope (CLSM) (Fluoview 3000, Olympus). The images were acquired using either 20x (Olympus, UPLFLN20XPH, NA=0.5) or 30x (Olympus, UPLSAP030XS, NA=1.05) objective, depending on the experiment. Acquired images were analysed using Fiji [48] and custom-made Python scripts. To assess the length of axon growth in the microchannels, images were overexposed and the length of the segmented line drawn on top of the axon was measured. See supplementary figure 2 for details.

### 2.5 Electrophysiology

6-channel and 60-channel spheroid-seeding PDMS microstructures were designed to fit the electrode layout of glass MEAs that were used to performed the electrophysiology experiments. During recording sessions, neurons cultured on MEAs were taken out of the incubator and placed in the MEA headstage (MEA2100-Systems, Multi Channel Systems) and kept at 5% CO_2_ without humidity control during 10 min recordings. The recordings were taken after four weeks in culture and each MEA was also imaged once at the end to assess the axon outgrowth.

The collected data sampled at 25 kHz was first filtered using a butterworth high pass filter with a cutoff frequency of 200 Hz. Spike detection was performed based on negative spike peaks. The baseline noise of the signal was calculated using the median absolute deviation (MAD). The peaks were considered spikes if their amplitude exceeded the baseline noise value 5 times. Additional peaks occurring within 3 ms after the first detected peak were discarded to avoid duplicates.

Spike-Triggered Time Histograms (STTHs) show the distribution of spike time latency between two electrodes. To obtain these distributions, spikes following in the first 16 ms after upstream-electrode spikes were used and binned into 0.05 ms sized time windows. To calculate the conduction speed, the distance between the respective electrodes was divided with a latency of the distribution peak:

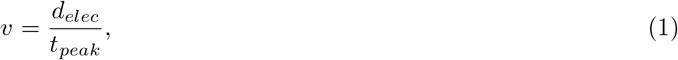

where *d*_*elec*_ in mm denotes the distance between the two respective electrodes, and *t*_*peak*_ in ms denotes the lower value of the time bin that contains a peak value of the latency distribution. Example of an STTH can be found in Supplementary Fig.5. In all electrophysiology data analyses, statistical significance was assessed using the Kruskal-Wallis test.

## 3 Results and discussion

In our investigation of how microstructure design impacts axon growth two similar, but distinct, experimental paradigms were used. For probing the impact of channel count and width we used spheroid cultures to ensure both the formation of axon bundles and to have an excess of axons compared to channels. In experiments investigating growth cone penetration through narrow restrictions this was unnecessary and as such low density cultures and associated microstructures (see section 2.1.2) were used.

### 3.1 Axon growth in spheroid-seeding microstructures with variable microchannel width

We seeded cell spheroids prepared as described in section 2.3.3 consisting of 500 neurons in the central seeding well of the PDMS microstructure. Emerging from the central well, microchannels initially narrow down to 1.5 *µ*m (Fig.2B) to ensure equal probability of similar number of axons entering each microchannel and then expand to the final microchannel width which increased logarithmically from 1.5 to 75 *µ*m (Fig.2A). We chose these widths because we believed that differences would be more prominent for lower channel widths (below 10 *µ*m). As seeding large neural spheroids provided an excess of axons, most channels were filled with axons (see S1 for confirmation).

Growth of GFP-labeled axons was assessed by imaging at three time points across three weeks *in vitro*. We imaged the samples at DIV 7, 12 and 23 and measured the extent of the axon length starting from the end of the narrowing/beginning of the final channel (see also Supplementary Fig.2). Since, we uniformly coated the dishes with PDL prior to mounting the microstructures, we expect equal density of axon guiding cues on the bottom of all microchannels, which implies that all putative differences arise from the differences in width of the PDMS microchannels. We observe a steep increase of axon length for microchannel widths up to 2.8 *µ*m (Fig 2C,D) after which there is a plateau at around 2 mm in length with a high variability at DIV7 (see also Supplementary Fig.4). A similar plateau is observed at 3 mm in length at DIV12 and 23. The growth of axons in topological constraint *in vitro* in two dimensional stiff substrate presents a significant difference when compared with axon lengths measured *in vivo*. Namely, though the variability is high, thalamocortical axon fibers in living rats reach total length of more than 6 mm *µ*m[11], which raises questions on the need for extracellular matrix that would provide three dimensional axon projections *in vitro* and thus increase the physological relevance of the *in vitro* platforms.

Notably, axons grow fastest in the first week of culture where projections in microchannels wider than 2.8 *µ*m reach roughly 2 mm in length. Between DIV 7 and DIV 12 the length revolves around 3 mm and it remains at this value at DIV 23. From this we conclude that in the absence of synaptic cues, the majority of axon growth happens withing the first two weeks in culture, which is consistent with former findings in the context of synapse formation[49], neural activity [50], and neurite growth[51]. Furthermore, it is worth discussing the profile of axons growing in microchannels. On Figure 2C we notice that axons in all channels tend to grow along the edge of the PDMS wall, which has already been reported elsewhere [40]. Channels smaller than 11.6 *µ*m in width tend to contain bundled axons, while in the wider channels the bundles tend to split into two sub-bundles each following one of the PDMS side walls (see Fig.2C).

### 3.2 The number of bundles per microchannel effects axon growth and AP propagation

In the next set of experiments, we report culturing 500-neuron spheroids in spheroid-seeding PDMS microstructures with constant channel width of 50 *µ*m (Fig.3A). We vary the number of channels that emerge from the central well in order to influence the average size of axon bundle formed in the microchannels. We expect the largest axon bundles in the 6-channel microstructure, while the 60-channel microstructure should contain the fewest axons per microchannel, hence forming the smallest axon bundles on average. Using 500-neuron spheroids would theoretically limit the number of axons to less than 9 per channel in the 60-channel microstructure, but we do not expect all neurons to project axons outside the spheroids. The support of this claim can be found in the supplementary material (Figure S6). Cells were cultured in PDMS on glass substrates and imaged on DIV 23. In Fig.3B we observe a decreasing trend in length for the increasing microchannel number. In other words, the thicker axon bundles tend to grow further. This finding can be related to the role of the axon-axon interaction in axon growth [52, 53].

Another possible explanation is that in 60-channel microstructures, in which the few axons can explore the full 50 *µ*m width of the channel, the increased available space may lead to axons changing direction more often during growth instead of growing straight along the microchannel (and adjacent axons). This conjecture regarding the influence of spatial constraints can be further substantiated by analyzing the morphology of axon bundles within these microchannels. Specifically, in the case of 6-channel microstructures designed to accommodate larger axon bundles, axons uniformly occupy all available space, likely facilitated by the higher density of axons within the bundle (Fig.3C). Conversely, in 60-channel microstructures, axon growth patterns resemble those observed in the experiments introduced in Subsection 3.1. Namely, they tend to form sub-bundles and follow the edges of the surrounding PDMS walls. Lastly, similarly as in the case of the microstructure with different channel widths, we observe high variability of the axon length as implied by the broad Gaussian distribution, especially in the case of the 6-channel microstructures.

Next, we placed the microstructures with the smallest (6) and the largest (60) number of microchannels on top of the MEAs to study the electrophysiological properties of axon bundles consisting of different number of axons. We calculated the mean firing rate per channel and the AP conduction speed as described in section 2.5.

We aligned 6-channel and 60-channel microstructures on top of MEAs using a brightfield microscope to obtain as many electrodes as possible per microchannel. This offered maximum 12 channel replicates per MEA. For a 60-electrode MEA of array shape 6×10 used in this experiments, the maximal number of electrodes per channel was five. We recorded spontaneous activity. In the case of 60-channel microstructure shown in Fig.4A, due to the limitation on the number of electrodes, only a selection of channels could be recorded, whereas the 6-channel microstructure (Fig.4B) was aligned on top of a MEA to obtain the recording from all 6 available channels. In Figure 4C we observe an increase of MFR per channel from week 2 to week 4 in culture. In fourth week in culture, we note a significant difference between MFRs for neurons in 6- and 60-channel microstructure respectively. In 60-channel microstructures, where there is presumably a lower density of axons per channel, we observe a higher MFR. This phenomenon can be attributed to the reduced number of axons within the microchannel, potentially enabling some of them to attain larger diameters within the same microchannel cross-section. Consequently, axon potentials of these axons generate larger extracellular potentials, rendering them more readily detectable by the electrode. There is another hypothesis stemming from our findings if we consider previously stated assumption that a lower number of microchannels could results in a higher density of axons per microchannel. This denser arrangement might limit the diffusion through the channels by increasing the tortuosity of the channels, thereby slowing down glucose and nutrient exchange and potentially leading to reduced neuronal activity [42].

**Fig. 4:**
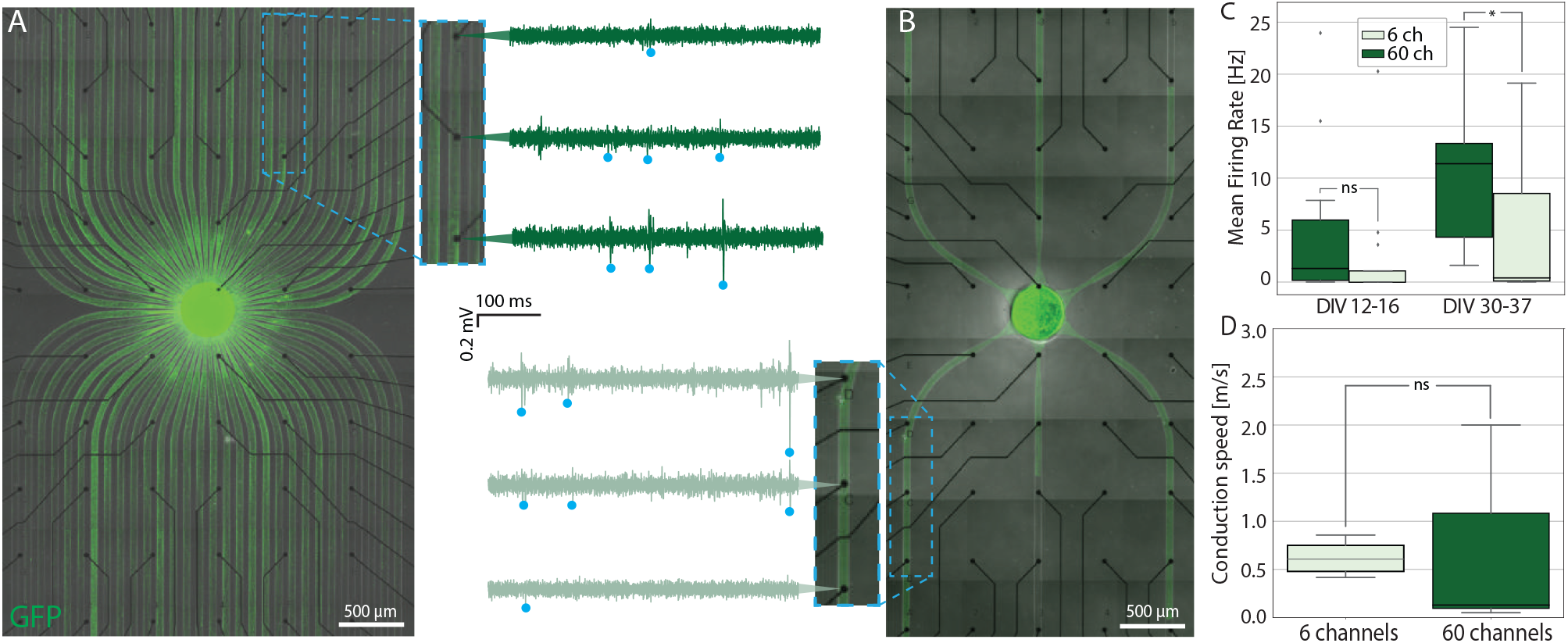
Spheroid-seeding PDMS microstructures for studying the electrophysiology of bundled axons. A) 60-channel PDMS microstructure aligned on top of a transparent 6×10 MEA and the corresponding voltage traces recorded from three selected electrodes along the same channel. B) 6-channel PDMS microstructure on top of a MEA and the corresponding voltage traces recorded from electrodes along the same channel. C) Mean firing rate recorded and calculated in the DIV range 12-16 and 30-37 for the cells in 6- and 60-channel PDMS microstructure respectively. A star corresponds to a p-value ¡ 0.05. D) AP conduction speed calculated using the delay times derived from spike-time triggered histograms in 6- and 60-channel PDMS microstructures. Number of biological replicates is 2.

In Fig 4D we calculated AP conduction speed for axons confined in 6- and 60-channel microstructure respectively for 4-week old cultures. Although there is no significant difference between them, we observe that the mean conduction speed for axon bundles within a 60-channel microstructure is lower, though the variability is higher. The higher variability in conduction speed for 60-channel microstructures can also be attributed to potentially having less axons per microchannel. Namely, as stated before, the excess of space allows for some axons to grow larger in diameter. Since the conduction speed depends also on axon diameter, higher variability in diameter could cause higher variability in conduction speed.

### 3.3 Growth cones able to penetrate 600 nm wide tunnels

In our subsequent experiments, we further explored the spatial confinement of axons using PDMS microchannels. Our aim was to probe the boundaries of axon growth and the flexibility of growth cones. Specifically, we sought to determine the minimal width of confinement required for a growth cone to adjust its size and traverse through. In order to eliminate the influence of other axons, we opted to seed cells in low-density suspensions.

In the first set of experiments, we designed a microstructure that consists of eight separate seeding wells that narrow down to a 30 *µ*m wide and 2 *µ*m high microchannel to ensure the axon guidance towards the microchannel (see Fig.5A). At the beginning of each microchannel there is an additional PDMS layer that narrows the available vertical space for axon growth to 600 nm. This narrowing we refer to as a submicron tunnel. Tunnels vary from 0.6 to 1.8 *µ*m in width presenting a spatial limitation in both horizontal and vertical direction. The last, eighth well does not begin with a tunnel but serves as a control channel of the axon growth in general.

**Fig. 5:**
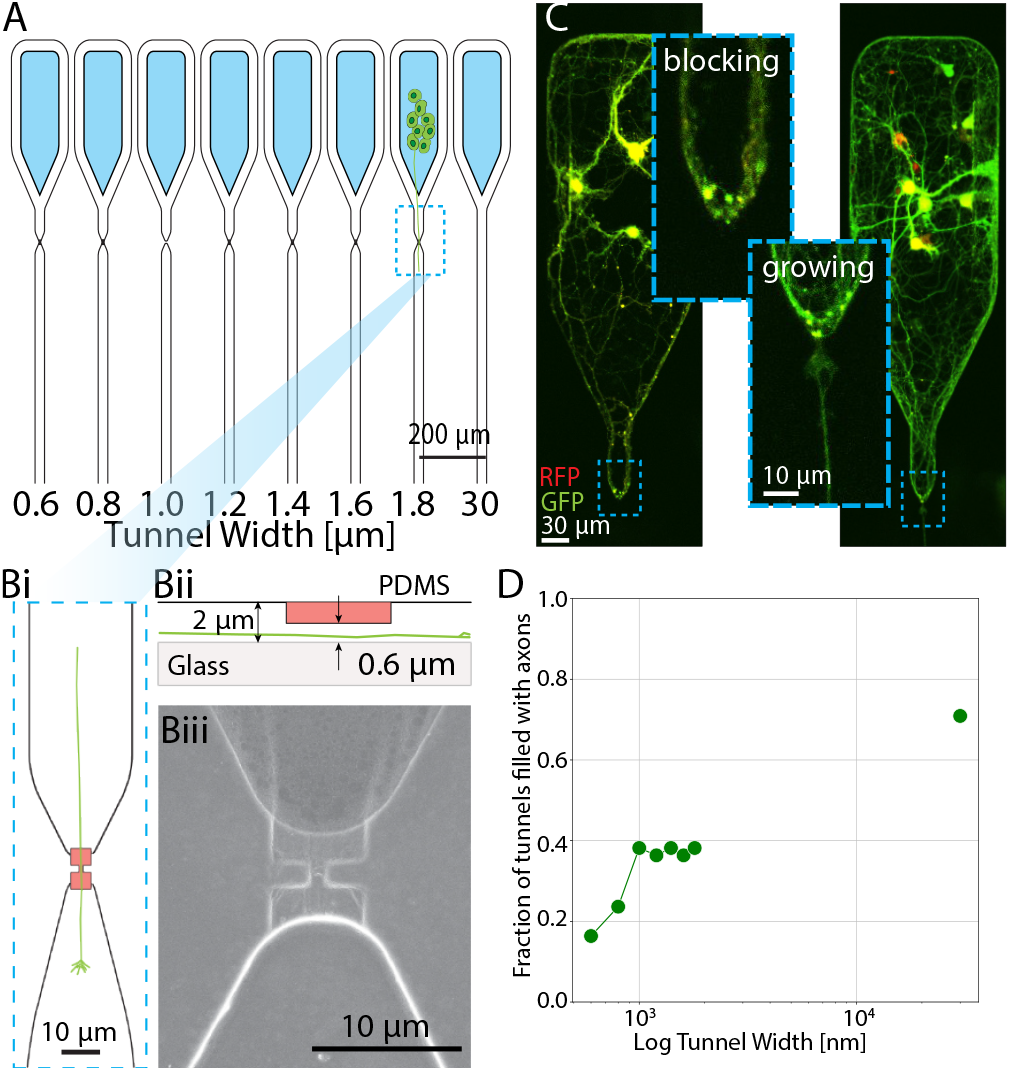
PDMS microstructures for studying axon growth through confinement. A) Neurons seeded into the seeding wells (blue) extend their axons towards the submicron-sized tunnel guided by narrowing microstructures. The tunnel width varies from 600 to 1800 nm while the subsequent microchannels have a fixed 30 *µ*m width. Bi) Schematic of the submicron tunnel region. Ideally axons grow directly through the narrowing. Bii) The tunnels are 600 nm high, while the rest of the channel has a height of 2 *µ*m. Biii) SEM image of submicron narrowing. C) Examples of axons turning and blocking the tunnel on the left and passing through the tunnel on the right. D) Fraction of tunnels that have axons passing through. There are 6 biological replicates.

Instances depicted in Fig.5C on the left, where axons curved and obstructed tunnel entrances, were noted. On the right of Fig.5C, we can observe an instance of an axon bundle narrowing down to successfully pass the tunnel. We observed that tunnels with widths equal to or larger than 1 *µ*m reached a plateau, with approximately 40% of these tunnels being filled (See Fig.5D). Our findings suggest that even with openings up to 1.8 x 0.6 µm^2^, neurons did not exhibit consistent growth. The impaired growth potential is an important aspect to be considered when designing microfluidic devices or an experimental setup in neural applications since it can affect the development of the cultured cells. If consistent growth is desired, both the narrowest point of the design and the channel width are important factors, as indicated by the results presented in this section and Section 3.1.

In the context of growth cone flexibility and the constraints on axon growth, we found that a width of 600×600 nm^2^ still allows axons to penetrate through in about 18% of our experiments. Considering that the average axon diameter is approximately 450 nm [54], and acknowledging the flexibility and adaptability of growth cones to their environment, this outcome is consistent with expectations.

### 3.4 Growth cones cannot penetrate tunnels narrower than 350 nm

Since in the previous section we have shown that axons can transverse through constrictions as small as 600 nm, in the last set of experiments, we decided to explore the very limitations of axon growth. We used the microstructure consisting of hexagonal seeding wells that narrow down to a 50 *µ*m wide and 2 *µ*m high area leading to a series of submicron tunnels that are 600 nm high (depicted in red in the zoomed in region of Fig.6A). With this microstructure we investigated tunnel widths varying from 100 to 1000 nm. In the previous set of experiments we observed that excess cell material at the narrow crossing points often blocked the tunnel entrance, so we designed this microstructure such that multiple tunnels share the same, wider microchannel. The open node transitions to 2 *µ*m high area that narrows down to a 600 nm area, which contains the submicron tunnel. Hence, each tunnel is 600 nm high but varies in width in the aforementioned range (100 to 1000 nm).

**Fig. 6:**
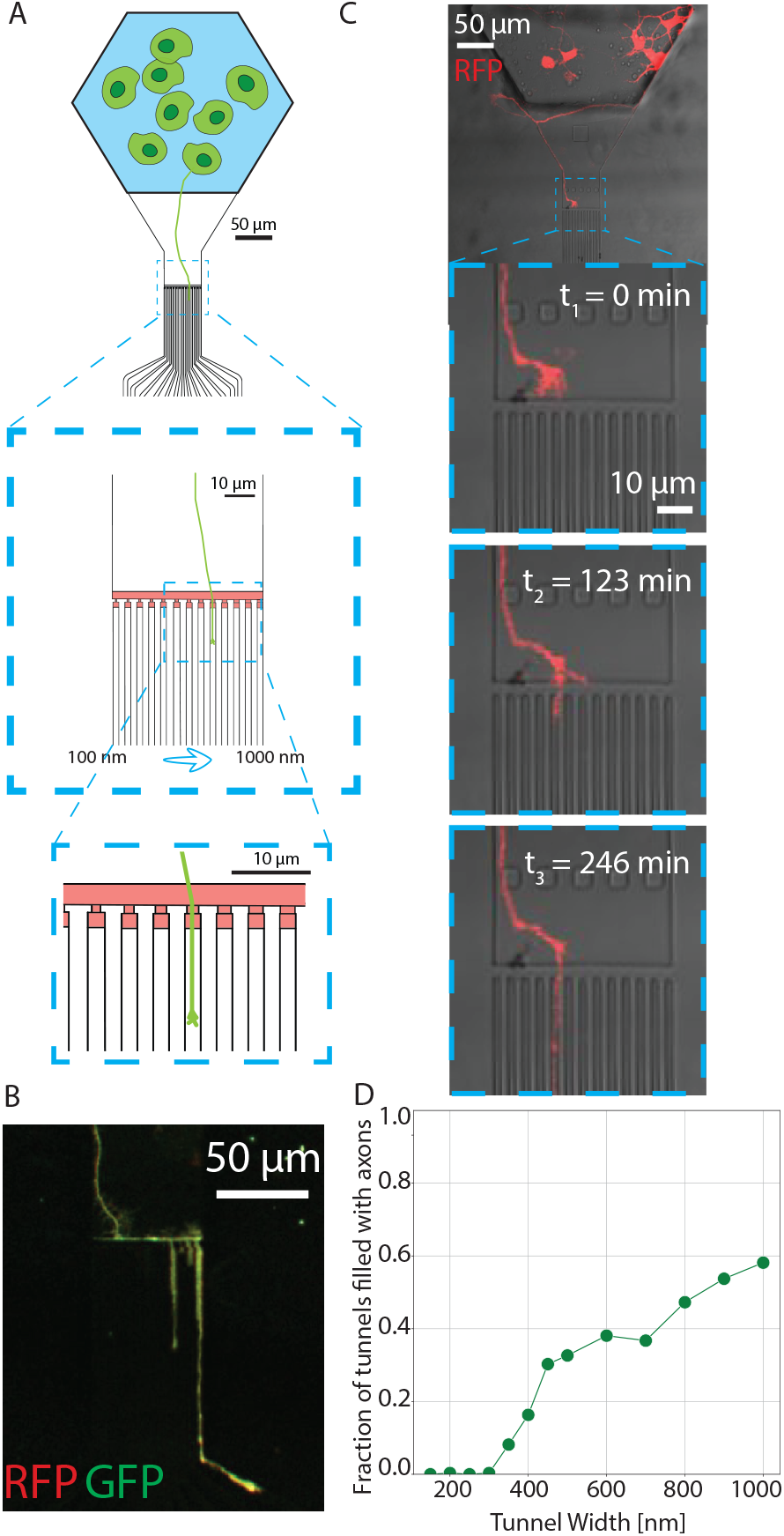
PDMS microstructures with smaller tunnels for studying the spatial limitations of axon growth. A) Microstructure schematic. Cells seeded in suspension fall into a hexagonal well. The axons are guided towards a submicron tunnel area and penetrate through. B) Example of a single axon branching through multiple tunnels. C) Overnight time lapse at DIV 2. D) Fraction of tunnels filled with channels. There are 6 biological replicates.

To investigate if growth cones are able to penetrate the tunnels, we imaged the cultures in DIV 10. We also performed an overnight time lapse on a subset of cultures on DIV 2 and DIV 3 to track axon movement and growth cone flexibility (full video is available in the supplementary material). In Fig.6B we observe a putative single axon growing along the tunnel area and splitting to penetrate three tunnels. The snapshots of the overnight time lapse video shown in Fig.6C were taken at *t*_1_ = 0 min, *t*_2_ = 123 min and *t*_3_ = 246 min. We observe the growth cone of approximately 8 *µ*m in diameter conforming its shape to fit into the 500 nm wide and 600 nm high tunnel. We notice that the same amount of time was needed for the growth cone to penetrate the tunnel and for the axon to afterwards grow for approximately 50 *µ*m. This is consistent with previous research that implies an increase of growth velocity in a topological constraint versus planar surfaces[55].

We did not observe any axons penetrating tunnels smaller than 350 nm. For wider channels we observe a gradual increase with channel width to reach the value 50% of channels with axons at 1 *µ*m, which is slightly higher than what we observed in the experiments described in Fig.5D. This could be due to the mentioned smaller likelihood of channels getting blocked by cell material with this design. This observation indicates that axons are more likely to enter wider microchannels. Similar overview of the effect of topographical constraints on axon growth has been shown[43]. These results further highlight the importance of micro-/submicro-environmental factors on axon growth.

## 4 Conclusions

We presented a comprehensive study focusing on axon growth within a spatially constrained *in vitro* environment. Our investigation demonstrated that the patterning of neuronal networks using PDMS microstructures offers a versatile platform for influencing axon growth, while allowing for both imaging and electrophysiology measurements. This platform provided a unique opportunity for long-term culturing of neurons in a controlled environment, enabling systematic exploration of parameter dependencies of the extent and limitations of axon growth, as well as changes in conduction speed. Notably, we maintained neurons in culture for over 4 weeks (we have recorded activity at DIV 37), and our colleagues and others have reported culturing neurons in similar microstructures for longer [35, 42].

The platform’s ability to immobilize neurons in one place facilitates systematic tracking of axon development *via* imaging. In addition, its compatibility with MEA allows for the assessment of action potential propagation. For the first time, we provided detailed insights into how spatial constraints influence axon growth. This knowledge is particularly significant in the context of advancing neural implants, especially living biohybrid neural implants designed to interface damaged tissue with artificially grown living replacements [56].

Furthermore, we identified areas for potential improvement in this system. Enhancing the data analysis pipeline to enable fast and reliable analysis of experimental data would eliminate the need for manual image analysis. Employing techniques such as atomic force microscopy combined with fluidics [57] would afford greater control over the size of neural networks and axon bundles forming in the microchannels. Integrating hydrogels into the system would allow exploration of axon behavior in 3D environments, enabling comparisons with electrophysiologically relevant or *in vivo* scenarios. Expanding the platform to isolate single cells would represent a significant advancement, facilitating full control and systematic analysis.

In summary, this platform holds promise for advancing both fundamental neuroscience research and translational applications, offering a versatile and adaptable framework for studying axon growth in controlled microenvironments.

## Author Contributions

KV: Conceptualization, Methodology, Software, Validation, Formal analysis, Investigation, Writing - Original Draft, Visualization. GA: Conceptualization, Methodology, Software, Investigation, Writing - Review & Editing. TR: Conceptualization, Methodology, Validation, Investigation, Data Curation, Writing - Review & Editing, Supervision, Project Administration, Funding Acquisition. RK: Investigation. SI: Conceptualization, Methodology, Software. JK: Software. JV: Conceptualization, Writing - Review & Editing, Resources, Supervision, Project Administration, Funding Acquisition. SW: Conceptualization, Methodology, Software, Validation, Investigation, Writing - Review & Editing, Visualization, Supervision, Project Administration.

## Conflicts of interest

There are no conflicts to declare.

## Acknowledgements

The authors would acknowledge ETH Zurich and Swiss National Science Foundation (SNSF) and the Human Frontiers Science Program (HFSP) for funding. The authors gratefully acknowledge Julian Hengsteler for SEM imaging. Authors are thankful to ScopeM for providing access to their SEM facility.

## Notes

### Competing Interest Statement

The authors have declared no competing interest.

