## Supplemental Figures for "Impact of microchannel width on axons for brain-on-chip applications"

### Supplementary material

In the Supplementary Table 1, one can find the list of all the PDMS microchannel and and (sub)micron tunnel widths used in the experiments with variable channel width, *i.e.* experiments described in Fig. 3, 5, and 6. For the experiments described in Fig. 3 and 4, microchannel size was constant, 50 $\mu$m with a microchannel height of 4 $\mu$m.

On Supplementary Fig. 1B) we can observe a pixel intensity profile along the arc that intersects the initial 1.5 $\mu$m narrowing of the microchannels, as shown on Supplementary Fig. 1A). The profile shows uniform distribution across all microchannels (Supplementary Fig. 1C)), implying the equal abundance of axons per channels. With this assumption, we can attribute all differences in the extent of axonal growth to topological constraints introduced by varying microchannel width.

To assess the extent of axon growth in the microchannels, image was log-processed in Fiji is ImageJ software for better visibility of flurosecence inside the channels. The same software was used to manually draw a segmented line along the channel until the fluorescence was visible by eye, as shown on Fig. 2. The length of this segmented line was used for further calculations.

Principle of two ways of cell seeding is shown on Supplementary Fig. 3.

On Supplementary Fig. 4 we show the mean length and the coefficient of variation for microchannels with variable channel width described in Fig. 2. Here we want to empahsize higher variability in length for smaller microchannels and additionally higher variability for earlier stages in culture.

An example of an STTH is shown on Supplementary Fig. 5 for three channels. The peak of the histogram is detected for every electrode and the distance between the corresponding electrode and the trigger electrode is divided by the time value of the lower edge of the peak time bin to get the conduction speed.

All microstructure types with GFP-expressing neural spheroids used in experiments described in Fig.3 and 4 are shown on Supplementary Fig.6A) and Fig.6C). All microstructures in the same row are seeded on the same dish and were imaged together, hence enabling the comparison of pixel intensity inside the channels. Average pixel intensity of a part of a channel was calculated for each microstructure. On Supplementary Fig.6B) and D) we see a decrease in intensity, implying the lower number of axons per channel for microstructures with higher number of channels.

Supplementary Fig.7 is the zoom out of the Fig.6 B), showing a neurite outgrowth inside the seeding well and branching of the axons in the submicron tunnels.

Table 1: List of channel and tunnel sizes for microstructures with variable width.

| Spheroid-seeding microstructure channel size | | Cell-suspension-seeding seeding microstructure channel size | | Cell-suspension-seeding seeding microstructure channel size | |
|---|---|---|---|---|---|
| Width [µm] | Height [µm] | Width [µm] | Height [µm] | Width [µm] | Height [µm] |
| 1.5 | 4 | 0.6 | 0.6 | 0.15 | 0.6 |
| 1.8 | 4 | 0.8 | 0.6 | 0.20 | 0.6 |
| 2.2 | 4 | 1.0 | 0.6 | 0.25 | 0.6 |
| 2.6 | 4 | 1.2 | 0.6 | 0.30 | 0.6 |
| 2.8 | 4 | 1.4 | 0.6 | 0.35 | 0.6 |
| 3.2 | 4 | 1.6 | 0.6 | 0.40 | 0.6 |
| 4.6 | 4 | 1.8 | 0.6 | 0.45 | 0.6 |
| 5.5 | 4 | 30.0 | 0.6 | 0.50 | 0.6 |
| 6.7 | 4 | | | 0.60 | 0.6 |
| 8 | 4 | | | 0.70 | 0.6 |
| 9.7 | 4 | | | 0.80 | 0.6 |
| 11.6 | 4 | | | 0.90 | 0.6 |
| 14 | 4 | | | 1.00 | 0.6 |
| 16.9 | 4 | | | | |
| 20.4 | 4 | | | | |
| 24.5 | 4 | | | | |
| 29.5 | 4 | | | | |
| 35.6 | 4 | | | | |
| 42.9 | 4 | | | | |
| 51.7 | 4 | | | | |
| 62.3 | 4 | | | | |
| 75 | 4 | | | | |

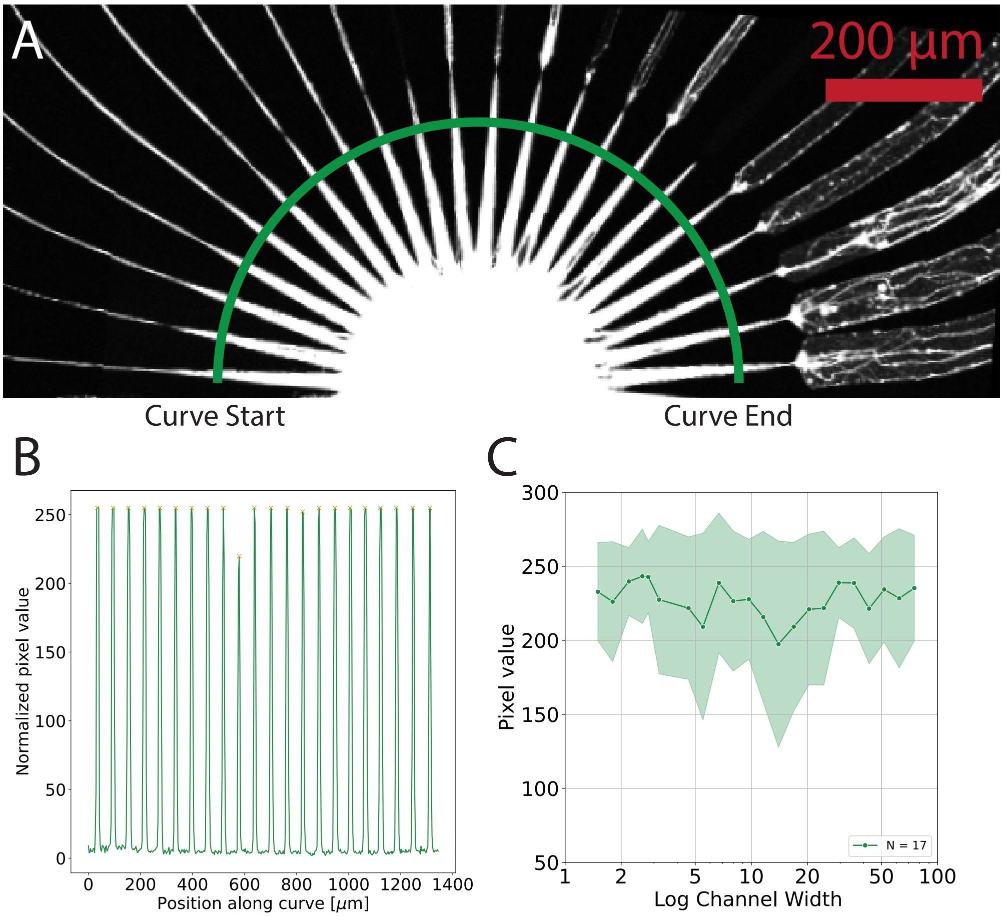


Figure 1: Spheroid-seeding microstructures with variable channel width. A) All channels are initially narrowed down to a 1.5$\mu$m wide channel to ensure equal distribution of axons. B) Intensity profile of a semi-circle intersecting the 1.5 $\mu$m microchannels. C) Average peak intensity per channel for 17 random samples.

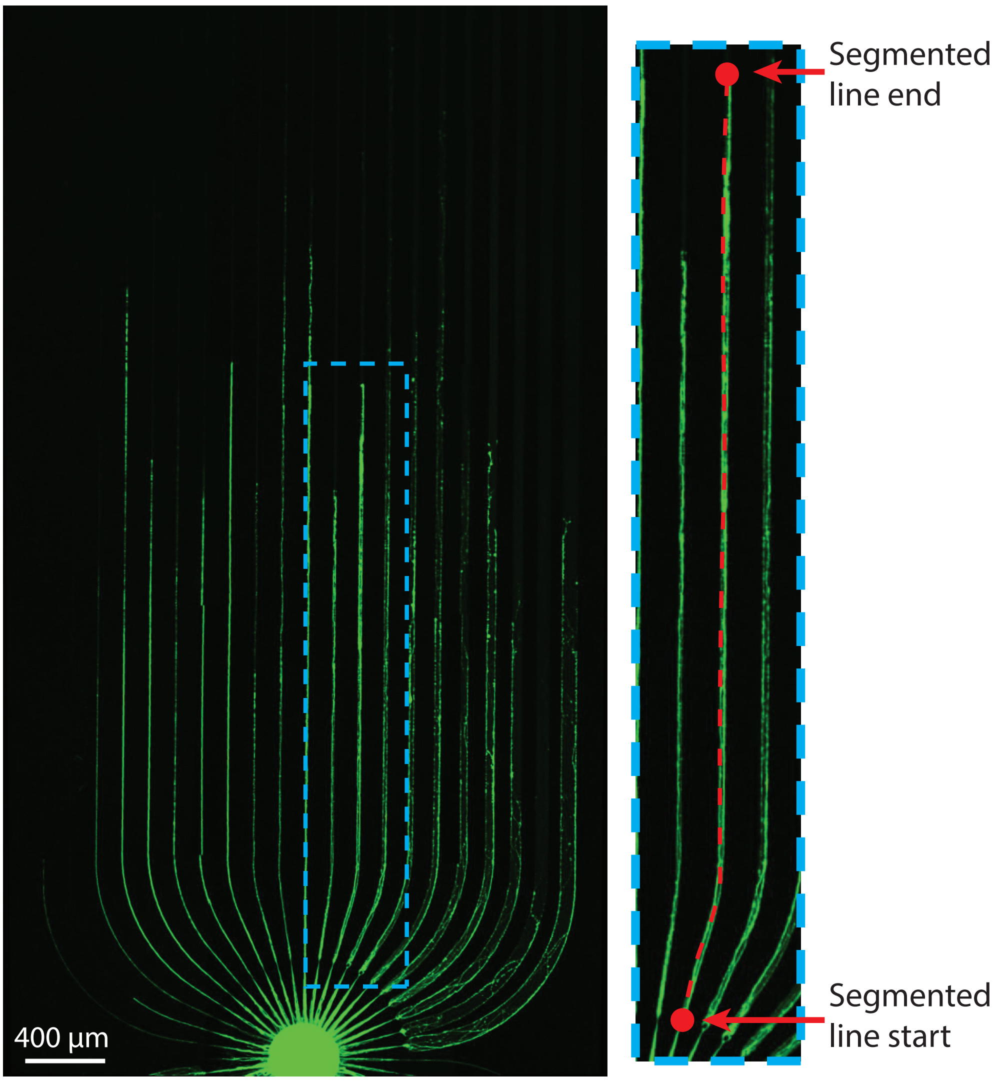


Figure 2: Analysis of the extent of the axon growth in spheroid seeding-microstructures. A segmented line was drawn and the length of the line was calculated using Fiji in ImageJ. The line begins at the end of the 1.5 $\mu$m microchannel and ends where there is no fluorescence anymore.

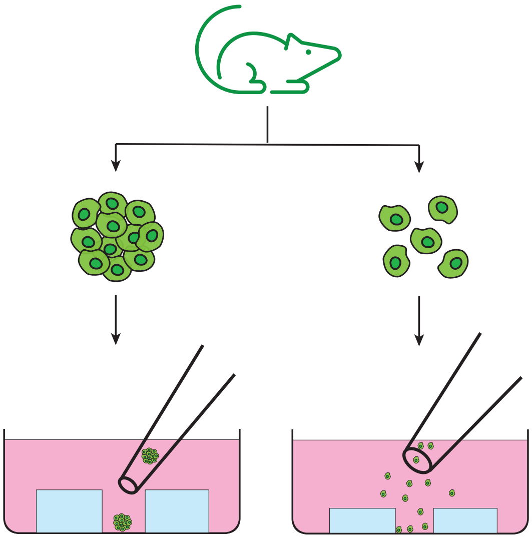

Figure 3: Two methods of seeding cells. Left: Spheroid seeding. 500-cell spheroids are pushed directly into the PDMS well using a 10 $\mu$m pipette. Right: Pipetting cell suspension above the PDMS using a 100 $\mu$m diameter glass pipette.

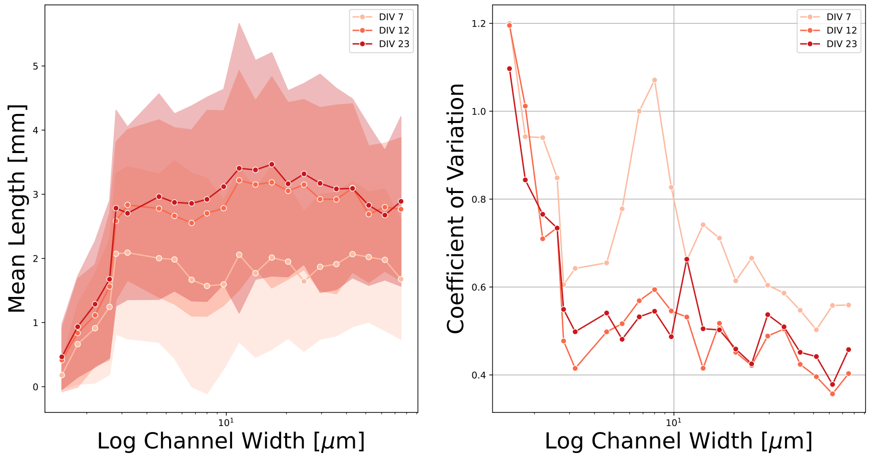


Figure 4: Mean and standard deviation of axon length in spheroid-seeding microstructures with variable microchannel width for different DIV.

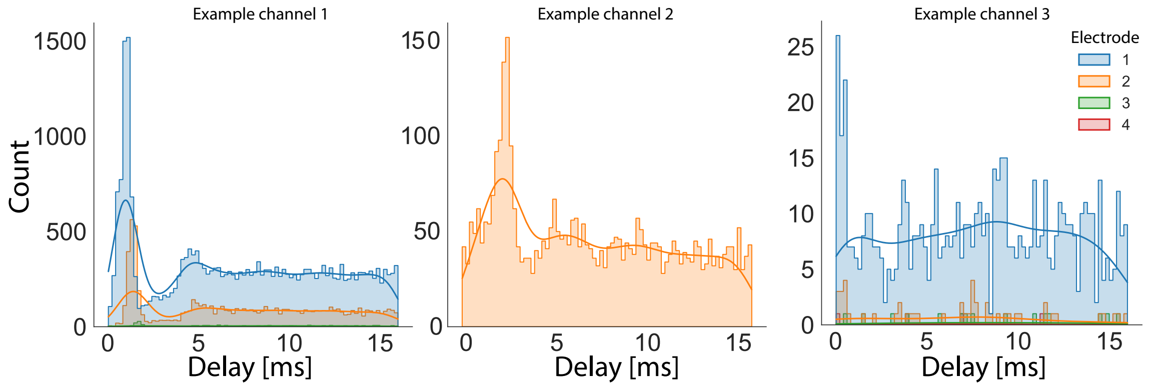


Figure 5: Spike time triggered histogram examples with the first electrode in the channel used as a trigger.

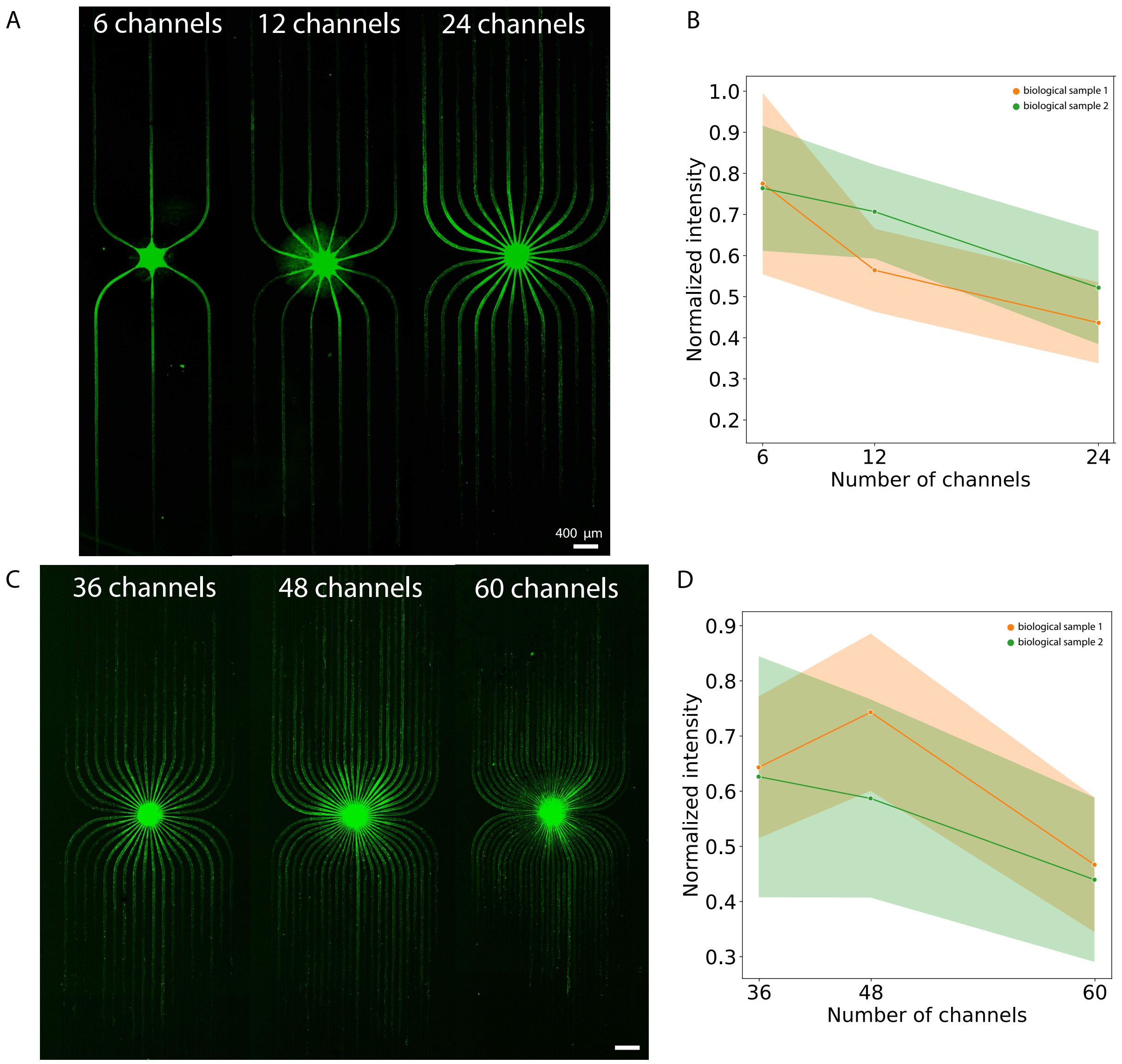


Figure 6: Examples of axon growth in spheorid-seeding microstructures with variable channel number. The pixel intensity within the channel was used as a metric for comparing the bundle size. We observe a decrease in mean pixel inensity for increasing number of channels, indicating a smaller bundle size when there are more microchannels emerging from the central well.

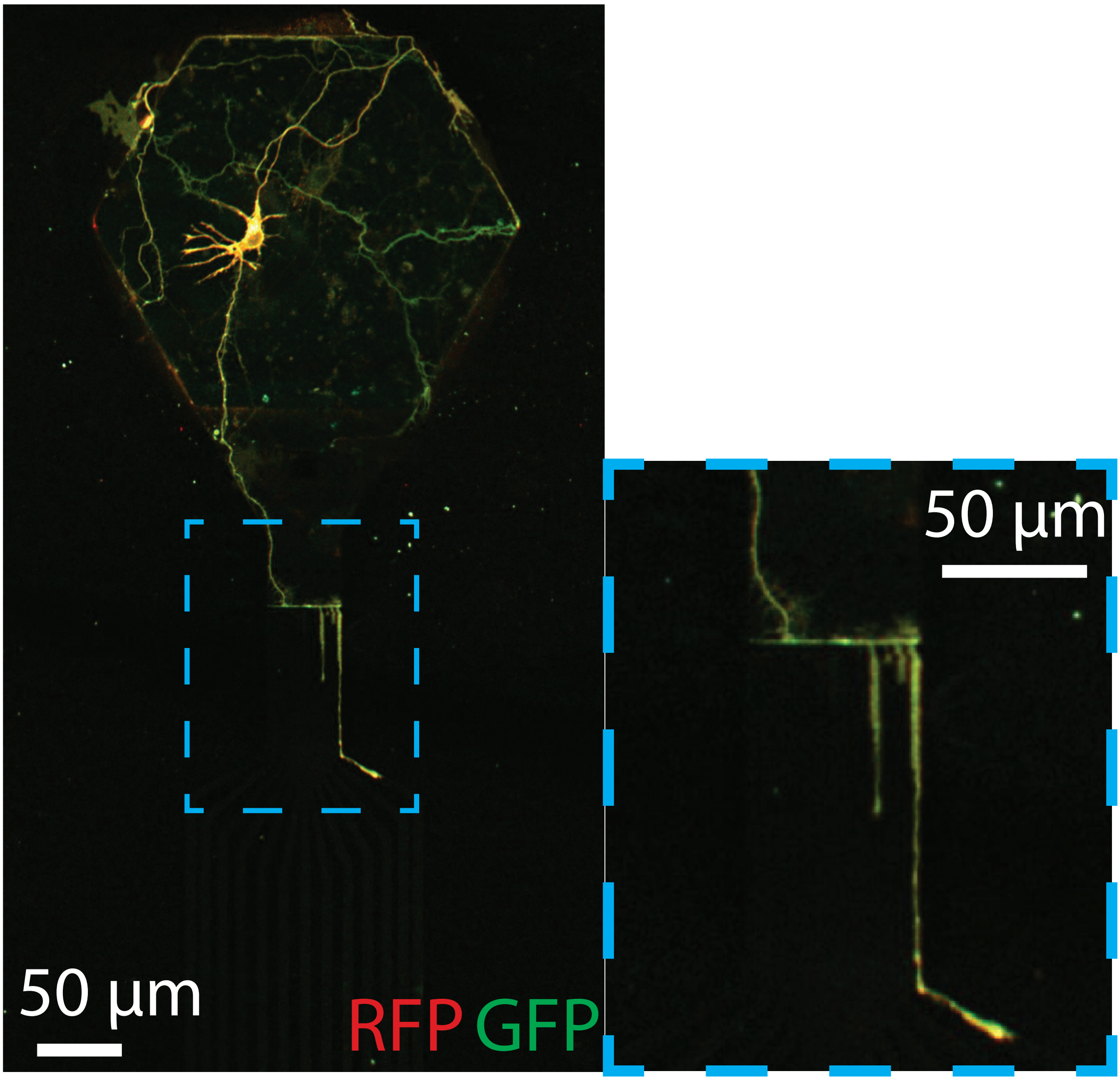


Figure 7: Overview of axon branching observed in the submicron tunnels shown in Figure 6 B).
